# *In vivo* production of pederin by labrenzin pathway expansion

**DOI:** 10.1101/2021.09.29.462382

**Authors:** Dina Kačar, Carmen Schleissner, Librada M. Cañedo, Pilar Rodríguez, Fernando de la Calle, Carmen Cuevas, Beatriz Galán, José Luis García

**Affiliations:** Department of Microbial and Plant Biotechnology, Centro de Investigaciones Biológicas, Agencia Estatal Consejo Superior de Investigaciones Científicas, Madrid, Spain; Research and Development Department, PharmaMar S.A., Madrid, Spain

## Abstract

Pederin is a potent polyketide toxin that causes severe skin lesions in humans after contact with insects of genus *Paederus*. Due to its potent anticancer activities, pederin family compounds have raised the interest of pharmaceutical industry. Despite extensive studies on the cluster of biosynthetic genes responsible for the production of pederin, it has not yet been possible to isolate and cultivate its bacterial endosymbiont producer. However, the marine bacterium *Labrenzia* sp. PHM005 was recently reported to produce labrenzin, the closest pederin analog. By cloning a synthetic *pedO* gene encoding one of the three *O*-methyltraferase of the pederin cluster into *Labrenzia* sp. PHM005 we have been able to produce pederin for the first time by fermentation in the new recombinant strain.

## Introduction

Pederin, a natural polyketide, has a huge therapeutic potential as a highly potent anticancer agent (Richter *et al*., 1997), however up to now the only source of this compound was the insect *Paederus fuscipes*, where it is found in very low amounts since twenty-five million field-collected insects had to be used to isolate the minimal amount of pure pederin (Pavan and Bo, 1952) to determine its chemical structure (Cardani *et al*., 1965). Pederin is produced by one of the first trans-AT mixed type polyketide/non-ribosomal peptide synthases (PKS/NRPS) assigned to a natural product that has been identified in an as-yet uncultivated bacterial symbiont of the insect (Piel, 2002; Piel, Höfer, *et al*., 2004; Piel, Wen, *et al*., 2004) and thus, its production by bacterial fermentation has not been achieved yet.

Pederin has among other structural peculiarities an *O*-methyl instead of the conventional ester or carboxylic acid polyketide terminus (Helfrich and Piel, 2010, 2016). In this sense, the cytotoxic activity of pederin-type compounds can be markedly increased by modifying the methylation pattern. The putative pederin biosynthetic gene cluster (*ped*) encodes three proteins with similarity to *O*-methyltransferases (MTs): PedA, PedE, and PedO (Piel, 2002; Piel, Hui, *et al*., 2004; Piel, Wen, *et al*., 2004). Biochemical *in vitro* experiments conducted with mycalamide as substrate, a pederin family compound with free C18-OH group, demonstrated that PedO was capable of introducing a methyl group at C18 position (Zimmermann *et al*., 2009).

Recently, the complete genome of the strain *Labrenzia* sp. PHM005, a free-living and cultivable producer of a pederin analog 18-*O*-demethyl pederin (Schleissner et al., 2017; Benítez *et al*., 2021), hereinafter labrenzin, has been sequenced (Kačar *et al*., 2019). A gene cluster responsible for the synthesis of labrenzin, named *lab* cluster, has been identified showing that it encodes a trans-AT mixed type PKS/NRPS biosynthetic pathway (Kačar *et al*., 2019). Interestingly, the *lab* cluster only encodes two MTs and this observation suggested that *Labrenzia* sp. PHM005 could be a good chassis to biosynthetically produce pederin by engineering the missing MT tailoring gene of *ped* cluster.

The aim of this work was to demonstrate that pederin could be produced by fermentation in *Labrenzia* sp. PHM005 by the heterologous expression of a *pedO* synthetic gene. In addition, further improvements of the pederin production have been investigated by overexpressing other MTs.

## Materials and methods

### Bacterial strains, media and growth conditions

Standard overnight *Escherichia coli* MFD*pir* (Ferrières *et al*., 2010) cultures were grown aerobically in Luria-Bertani (LB) broth or LB agar at 37 °C (Bertani, 1951). The medium was supplemented with 1 mM diamine-pymelic acid (DAP) and the corresponding antibiotic, when appropriate. *Labrenzia* sp. PHM005 wild type and recombinant strains were grown in Marine Broth (MB) Difco 2216 (Sigma-Aldrich) or Marine Agar (MA) Difco 2216 (Sigma-Aldrich), supplemented with antibiotics, when appropriate. All the strains were cultured in 50 mL falcon tubes or 100 mL flasks with 10 and 20 mL of medium, respectively. Culture medium used to study labrenzin and pederin production in *Labrenzia* sp. PHM005 was modified using marine basal medium supplemented with vitamins (MBM+vit) (Kačar *et al*., 2019). The culture medium was supplemented with 0.2 mM 3-methyl-benzoate, when the cloned gene was expressed under the control of the inducible *Pm* promoter. The strains were grown overnight in falcon tubes in MB at 30 °C with shaking at 200 rpm. The overnight culture was washed in 0.85% NaCl solution and diluted to an optical density (OD_600_) ≈ 0.1 in 20 mL of fresh medium. To determine the production of labrenzin, pederin and analogs by HPLC/MS analyses, the strains were cultured for 72 h.

### Plasmid DNA transformation and clone selection

*E. coli* MFD*pir* electro-competent cells were prepared and transformed by electroporation as described (Wirth *et al*., 1989). To select the transformants chloramphenicol (34 μg/mL) and 1 mM DAP was added to the LB agar plates. A biparental conjugation was used for transformation of *Labrenzia* sp. PHM005 using *E. coli* MFD*pir* carrying the plasmid of interest as a donor strain. Samples of 1 mL of overnight cultures of *Labrenzia* and *E. coli* MFD*pir* were collected by centrifugation and the pellets were washed with 500 μL of 0.85% NaCl and resuspended into 200 μL of the same solution. Samples of 50 μL of each strain were mixed and deposited into a 0.22 μL filter mating disc placed on the surface of a MA agar plate that was further incubated for 4-6 h at 37 °C. The cells deposited on the filter disc were collected with 1 mL of 0.85 % NaCl and vortexed thoroughly to detach the cells from the filter. Afterwards, cells were plated by dilutions on MA plates containing chloramphenicol (5 μg/mL).

### Construction of plasmids for expression of MTs

Plasmid pSEVA338 from pSEVA collection (http://seva-plasmids.com/) was used as vector to create an artificial operon containing three genes encoding the Lab6 and Lab16 MTs from *Labrenzia* and PedO MT from *Paederus* symbiont. The artificial operon was designed as indicated in Figure S1, where each gene with the corresponding RBS was flanked by specific restriction sites (blunt cut) in a way that individual genes or the combination of genes could be easily generated by digestion and re-ligation. The operon was synthetized and cloned using *Sac*I and *Spe*I restriction sites by GenScript yielding plasmid pSEVA338_MTs. Other plasmids derived from pSEVA338_MTs were pSEVA338_*lab6*, pSEVA338_*lab16*, pSEVA338_*pedO*, pSEVA338_*lab6*_*lab16*, pSEVA338_*lab6_pedO* and pSEVA338_*lab16*_*pedO*. The inducible promoter *Pm* from pSEVA338 was replaced by the strong *p14g* promoter in plasmid pSEVA227M (kind gift from Gonzalo Durante) using restriction enzymes *Pac*I and *Avr*II generating a new construct pSEVA338_*p14g*_*lab6*_*pedO*.

### Extraction, purification and identification of polyketide compounds

Upon fermentation 20 mL of culture medium collected by centrifugation was frozen at −80 °C and subsequently freeze dried. The lyophilized product was then dissolved in 4 mL of distilled water and equal volume of ethyl acetate. After homogenization and centrifugation, the organic phase was collected by pipetting and the extraction was repeated once more. The collected organic phase was dried by vacuum centrifugation and the pellet was dissolved in 150 μL of methanol and filtered for a HPLC/MS analysis.

HPLC-MS analysis was carried out using a HPLC/MS system and a separation column previously described (Kačar *et al*., 2019). The running method was as follows: solvent A was 100% water and solvent B was 100% acetonitrile. The flow rate was 500 μL min^-1^ using the following gradient: t = 2 min, 100% A; t = 8 min, 95% A; t = 40 min, 55% A; t = 53 min, 0% A; t = 55 min, 0% A; t = 57 min, 100% A; t = 65 min, 100% A.

## Results

### Genetic analyses

The unique structural difference between pederin and labrenzin is the absence of an *O*-methylation of C18-OH in labrenzin (Figure 1). The absence of such methylation can be justified assuming that the *lab* cluster contains only two MTs, *i.e*., Lab6 (MT6) and Lab16 (MT16) (Kčar *et al*., 2019), whereas *ped* cluster contains three MTs, *i.e*., PedA, PedE and PedO. A protein homology comparison between MTs from *lab* and *ped* clusters showed that Lab16 is homologous to PedE with a 51% amino acid sequence identity and Lab6 shares 47% sequence identity with PedA. According to Blast analysis the sequence amino acid identity of PedO with Lab16 was very low (28% with a query cover of 63%) but Lab6 and PedO showed a 53% sequence identity. Although the genetic analysis cannot exclude Lab6 from the experimental setting, the previous *in vitro* biochemical analysis carried out with PedO using mycalamide A as substrate demonstrated that PedO is responsible for the methylation of the C18-OH group in pederin (Zimmermann *et al*., 2009). Thus, we assumed that PedO could be the missing gene in *Labrenzia* responsible of the C18-OH methylation in the bacterial symbiont.

**Figure 1.**
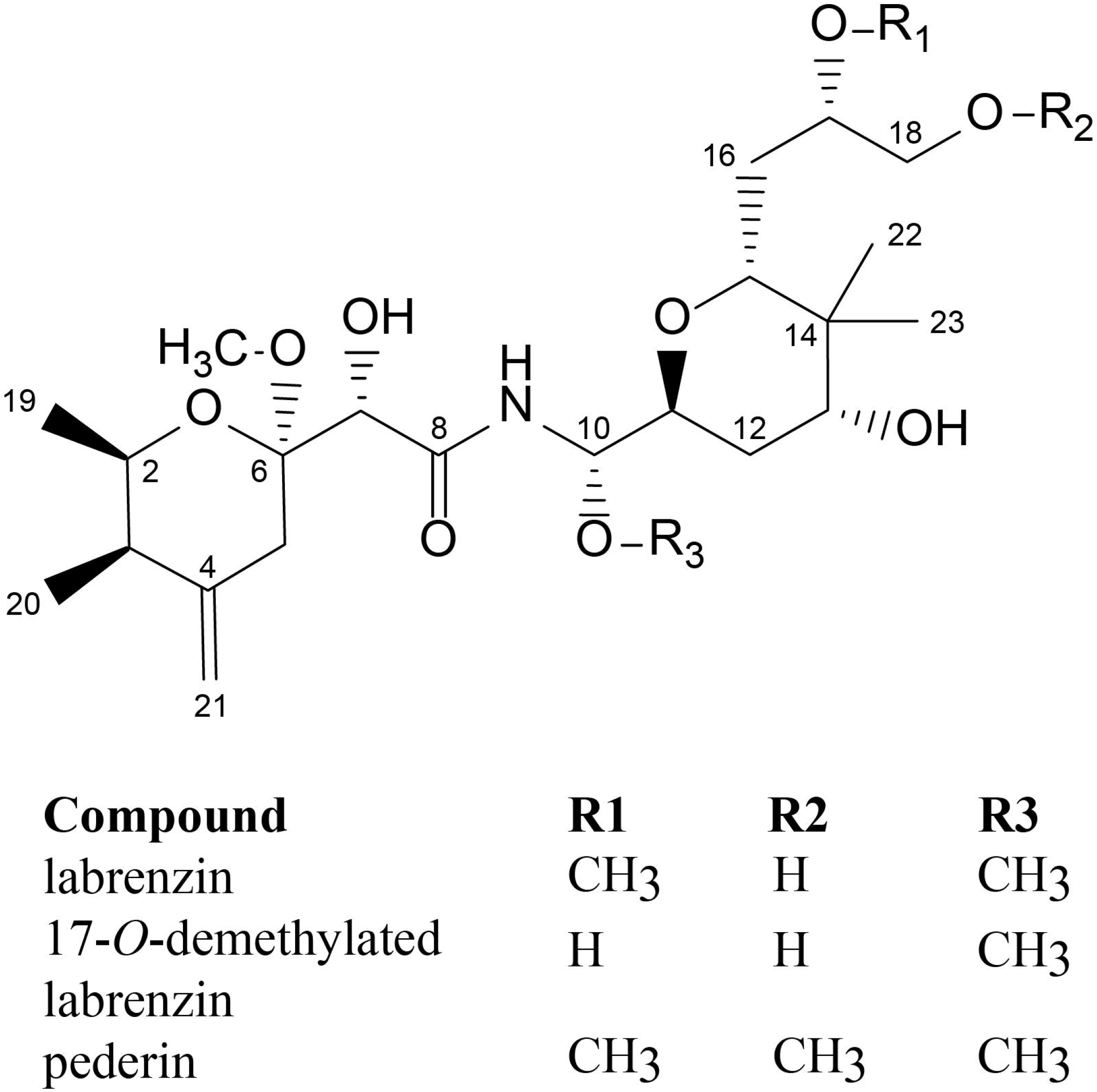
Chemical structure of pederin family polyketides.

Nevertheless, considering that PedO methylation activity was only tested *in vitro* using mycalamide as substrate our hypothesis required an experimental *in vivo* demonstration since PedO could not methylate labrenzin under the *in vivo* environmental conditions where the concentrations of the reaction substrates inside the cells are different than those used for the methylation of mycalamide *in vitro* and different labrenzin intermediates could compete as putative substrates or even work as inhibitors. Moreover, although PedO was produced in an active form in *Escherichia coli* after the optimization of its expression (Zimmermann *et al*., 2009), the gene and/or the enzyme could require some additional optimization to become active in *Labrenzia*.

### Production of pederin in Labrenzia sp. PHM005

To test our hypothesis, this is, to be able to synthetize pederin for the first time in a cultivable bacterium, a synthetic *pedO* gene was heterologously expressed in *Labrenzia* sp. PHM005 transformed with the recombinant plasmid pSEVA338_*pedO* harbouring the *pedO* synthetic gene under the control of the *P_m_* inducible promoter. As predicted, when a culture extract of *Labrenzia* sp. PHM005 (pSEVA338_*pedO*) was analyzed by HPLC/MS we observed a new intermediate, more hydrophobic than the previous pederin analogs, *i.e*., compound (1) (labrenzin) and compound (2) (17-*O*-demethylated labrenzin) produced by the wild type strain (Figure 2). The MS spectrum revealed a new peak (compound 3) observed at RT = 40.4 min and with ion fragmentation m/z= 526; 440; 422. This fragmentation pattern is matching with both labrenzin and 17-*O*-demethylated labrenzin (Figure S2) and indicates an additional methyl group (M+14), as it is in pederin.

**Figure 2.**
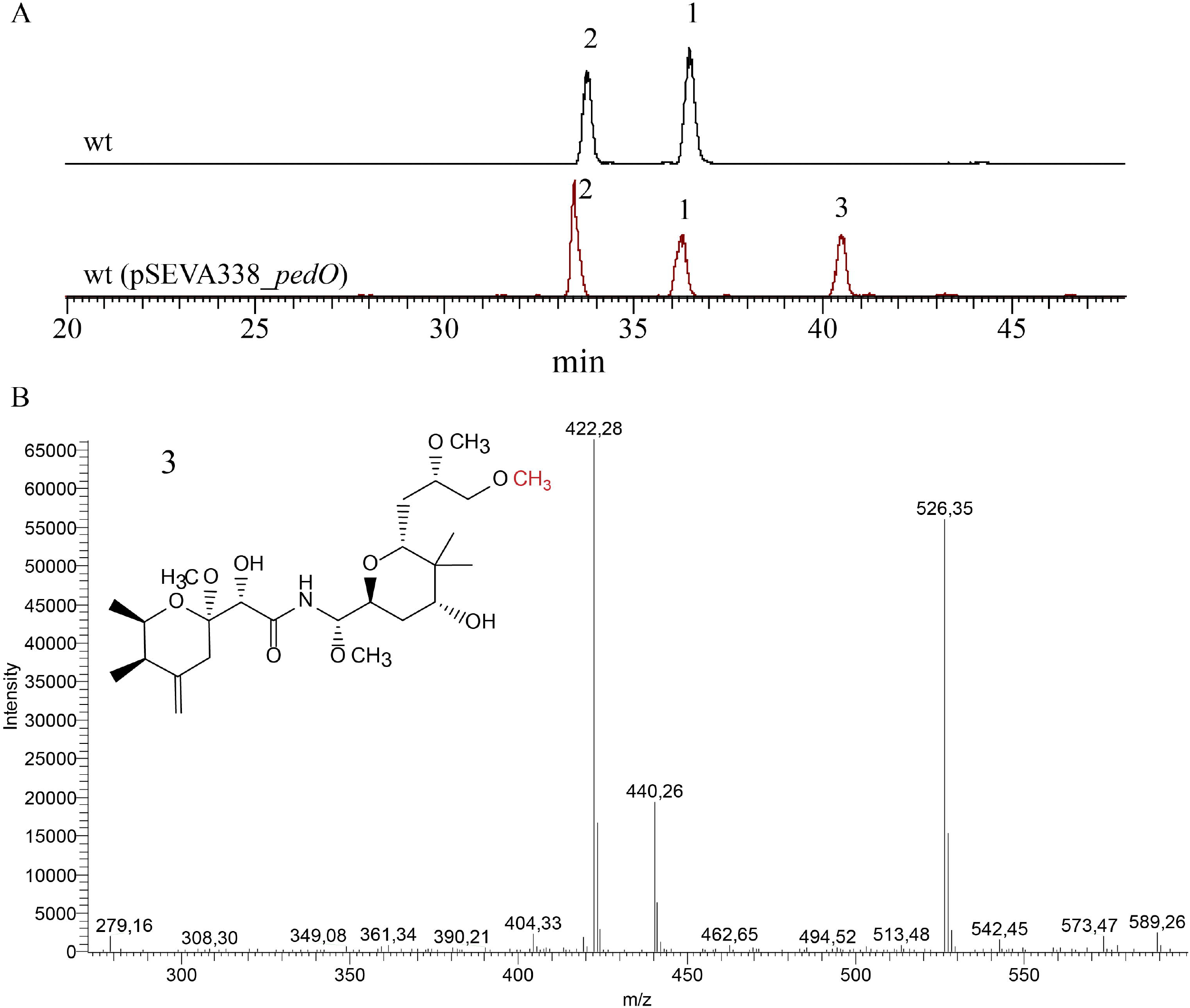
Pederin biosynthesis in *Labrenzia* sp. PHM005 wt (pSEVA338_*pedO*). A) HPLC-MS chromatograms of the supernatant extracts obtained after 72 h of cultivation of wt cultures in MBM+vit medium presenting extracted ions in the range m/z=498-526. Intermediates compound (1) (labrenzin), compound (2) (17-*O*-demethylated labrenzin) and compound (3) (pederin) are indicated. B) ESIMS ion fragmentation of compound (3) (pederin).

### Co-overexpression of lab6, lab16 and PedO methylases

Although pederin was produced in *Labrenzia* sp. PHM005 (pSEVA338_*pedO*), it was not the most abundant intermediate in the extract. Therefore, to optimize the pederin production we analysed the production in different strains co-expressing the *pedO* gene in different combinations with *lab6* and *lab16*. To this aim, plasmids pSEVA338_*lab6*_*pedO*, pSEVA338_*lab16*_*pedO*, pSEVA338_*lab6*_*lab16*_*pedO* were constructed and transformed in *Labrenzia* sp. PHM005. The resulting strains were cultivated in the production medium and the production of the labrenzin analogues was analysed. Figure 3 (A-E) shows HPLC-MS chromatograms of culture extracts of strains harnessing different expression combinations of MTs using pSEVA338 plasmid. The first conclusion was that the relative pederin amount was increased by co-expressing *pedO* with *lab6* and *lab16* genes in the different combinations tested (Figure 3E and 3F). 17-*O*-demethylated labrenzin (compound 2) was the most abundant peak when *pedO* was expressed alone forming an operon with *lab16* and when the three MTs encoding genes were co-expressed (Figure 3C and 3D). However, *pedO* overexpressed together with *lab6*, appeared to be the optimal combination for pederin production under the culture conditions.

**Figure 3.**
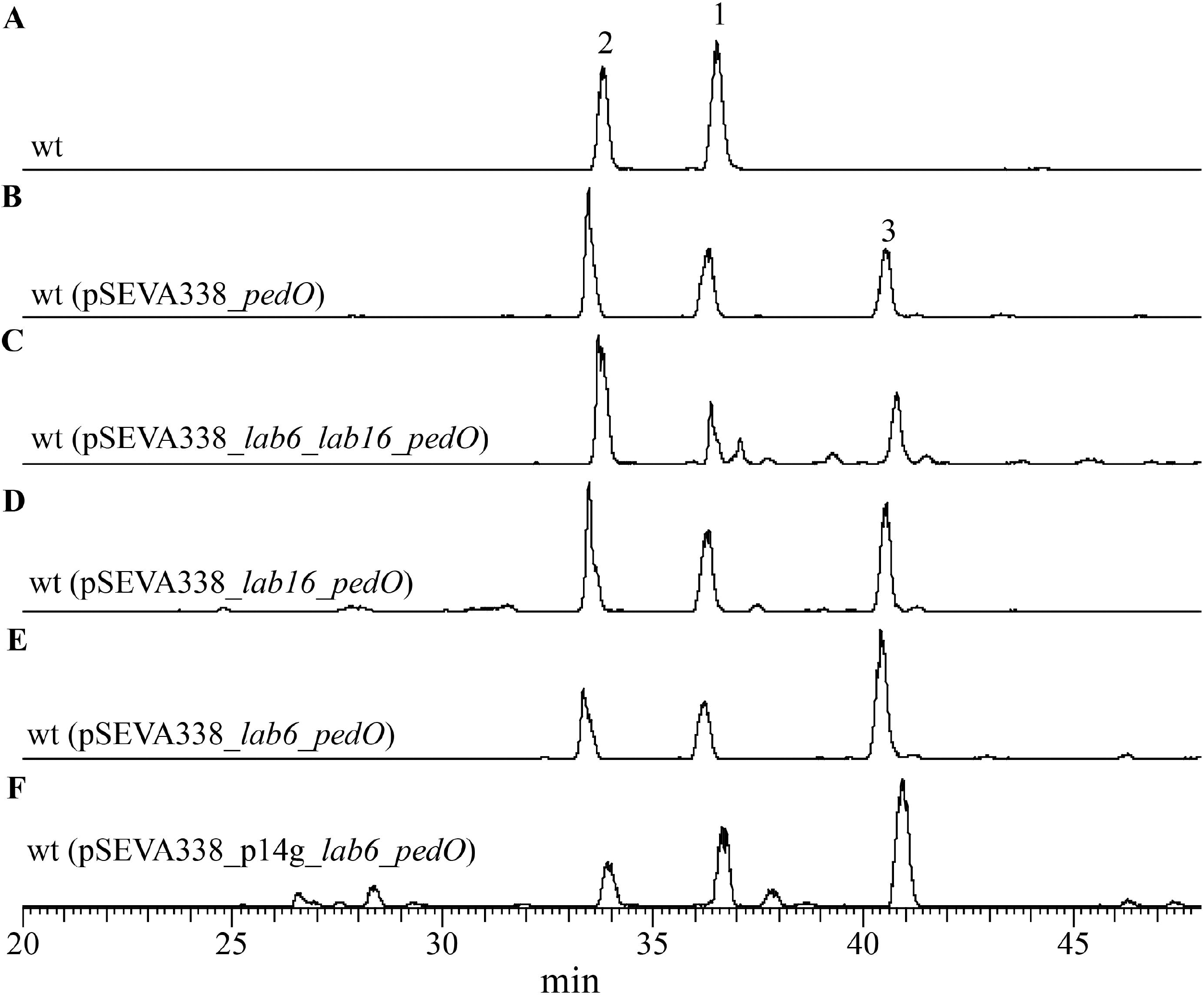
HPLC-MS chromatograms of the culture extracts obtained after 72 h of cultivation of wt recombinant strains transformed with different plasmids in MBM+vit medium. Extracted ions range is m/z=498-526. Intermediates 1 (m/z= 512), 2 (m/z= 498) and 3 (m/z= 526) are indicated. A) wt; B) wt(pSEVA338_*pedO*); C) wt(pSEVA338_*lab6*_*lab16*_*pedO*); D) wt(pSEVA338_*lab16_pedO*); E) wt(pSEVA338_*lab6_pedO*); F) wt(pSEVA338_p14g_*lab6_pedO*).

### Engineering the promoter driven the expression of the lab6_pedO MTs

The second approach was to engineer the expression plasmid pSEVA338_*lab6*_*pedO* with the constitutive strong *P_14g_* promoter, replacing the inducible *P_m_* promoter. Previously, we tested the *P_14g_* promoter strength in *Labrenzia* sp. PHM005 by constructing a transcriptional fusion with GFP showing that it is functional and has high expression levels (data not shown). The resulting plasmid named pSEVA338_*p14*_*lab6*_*pedO* was transformed in *Labrenzia* sp. PHM005. The production of the recombinant strain was analysed showing that the promoter switch did not seem to alter the peak ratio as observed in the MS chromatograms (Figure 3F), suggesting that the expression of *pedO* is not the bottleneck of the process.

## Discussion

*O*-Methylation modulates the pharmacokinetic and pharmacodynamic properties of natural products, affecting their bioavailability, stability, and binding to targets. Tailoring the polyketide structures allows an additional level of functional complexity, and thus, polyketide pathway engineering has generated new-to-nature products through novel glycosylation, acyltransferase, hydroxylation, epoxidation, alkylation, transamination and desaturation reactions acting on naturally occurring products (Cumming *et al*., 2014). However, as far as we know, very few experiments have been carried out to expand a polyketide pathway to generate novel polyketides by cloning tailoring *O*-MTs. Many years ago, Fu *et al*. (1996) expressed the TcmO *O*-MT of the tetracenomycin biosynthetic pathway of *Streptomyces glaucescens* was expressed in *Streptomyces coelicolor* CH999 together with the actinorhodin polyketide synthase (PKS) gene cluster, which is responsible for the biosynthesis of 3,8-dihydroxy-methylanthraquinone carboxylic acid (DMAC) and its decarboxylated analog, aloesaponarin. The resulting recombinant strain produced approximately equal quantities of aloesaponarin and a new product but no DMAC. More recently, Wang *et al*. (2019) have studied the use of two fungal MTs to produce unnatural O-methylated benzenediol lactone polyketides.

In this work, we have confirmed the role of PedO MT found in *Paederus* bacterial symbiont. In addition, we have demonstrated that its heterologous expression in recombinant strains of *Labrenzia* sp. PHM005 has allowed expanding the labrenzin biosynthetic pathway generating a new compound, pederin, providing labrenzin with an additional methylation on C18-OH. As mentioned above, pederin can be only isolated from beetle extraction so far, and thus, this is the first time pederin is produced by direct fermentation in a cultivable bacterium. Therefore, pederin could be now produced by fermentation at large scale to be tested and used as an antitumoral drug. Nevertheless, although we have developed different *pedO* expression systems for the production of pederin further improvements should be made for its efficient industrial scale production since the production levels of labrenzin and pederin are still low. In addition to increasing production of the labrenzin polyketide scaffold, we have shown that a finetuning of the expression of *lab6* and *pedO* genes should be optimized to increase the pederin production. It is possible that sequential *O*-methylation tailoring would depend on the substrate preference of MTs resulting the synthesis of the intermediates more favorable than that of pederin. A similar finding was observed when PedO was unable to methylate the C18-OH group of mycalamide A intermediate when the neighboring - OH was methylated, *i.e*., C18 *O*-methylation appeared only possible when the C17-OH was non methylated (Zimmermann *et al*., 2009). On the other hand, the synthetic *pedO* gene used in this study originates from an uncultured symbiont bacterium (Piel *et al*., 2005), therefore, its expression might not be very efficient in *Labrenzia* sp. PHM005. In that sense, for more efficient pederin transformation from labrenzin in the industrial setting, we suggest to carry out codon usage modifications or to approach a directed evolution of PedO enzyme. Finally, it is important to consider that the production of a combination of different sequentially methylated intermediates, all of them secreted to the surrounding medium, can provide an advantage for the producing bacteria, since different products can develop different roles in their surrounding environment. Thus, the simultaneous production and secretion of the intermediates can be detrimental for the industrial overproduction of pederin, since we have determined that labrenzin once secreted to the medium cannot be uptake by the cells (data not shown) and therefore, it cannot be methylated.

## Supporting information

Figure S1. Scheme representing the design of the artificial operon comprising PedO-Lab6-Lab16 methyltransferases, under the control of the Pm promoter

Figure S2. ESIMS ionization of A) labrenzin and B) 17-O-demethyl labrenzin.

## Acknowledgements

The present study was funded by the Ministry of Economy and Competitiveness of Spain under the program RETOS-COLABORACIÓN with the project number RTC-2016-4892-1 (DESPOL). The authors would like to thank Ana Valencia for technical assistance.

---

**Figure S1.** Scheme representing the design of the artificial operon comprising *PedO-Lab6-Lab16* methyltransferases, under the control of the *Pm* promoter of pSEVA338.

**Figure S2.** ESIMS ionization of A) labrenzin and B) 17-*O*-demethyl labrenzin.

