## Supplementary figures and images for "*In vivo* production of pederin by labrenzin pathway expansion"

### Figure S1. Scheme representing the design of the artificial operon comprising PedO-Lab6-Lab16 methyltransferases, under the control of the Pm promoter

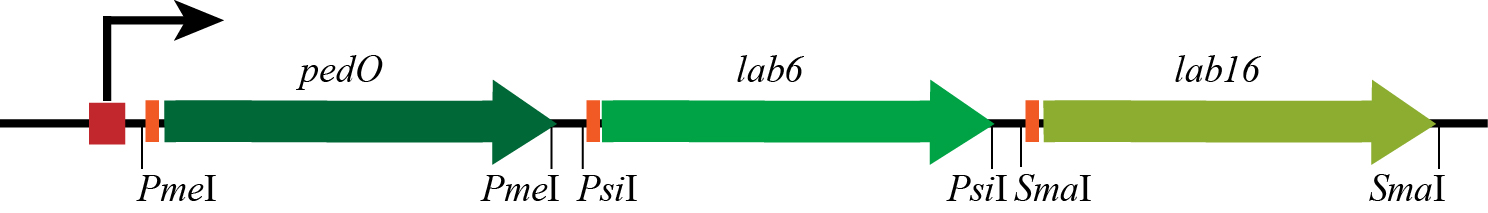

### Figure S2. ESIMS ionization of A) labrenzin and B) 17-O-demethyl labrenzin.

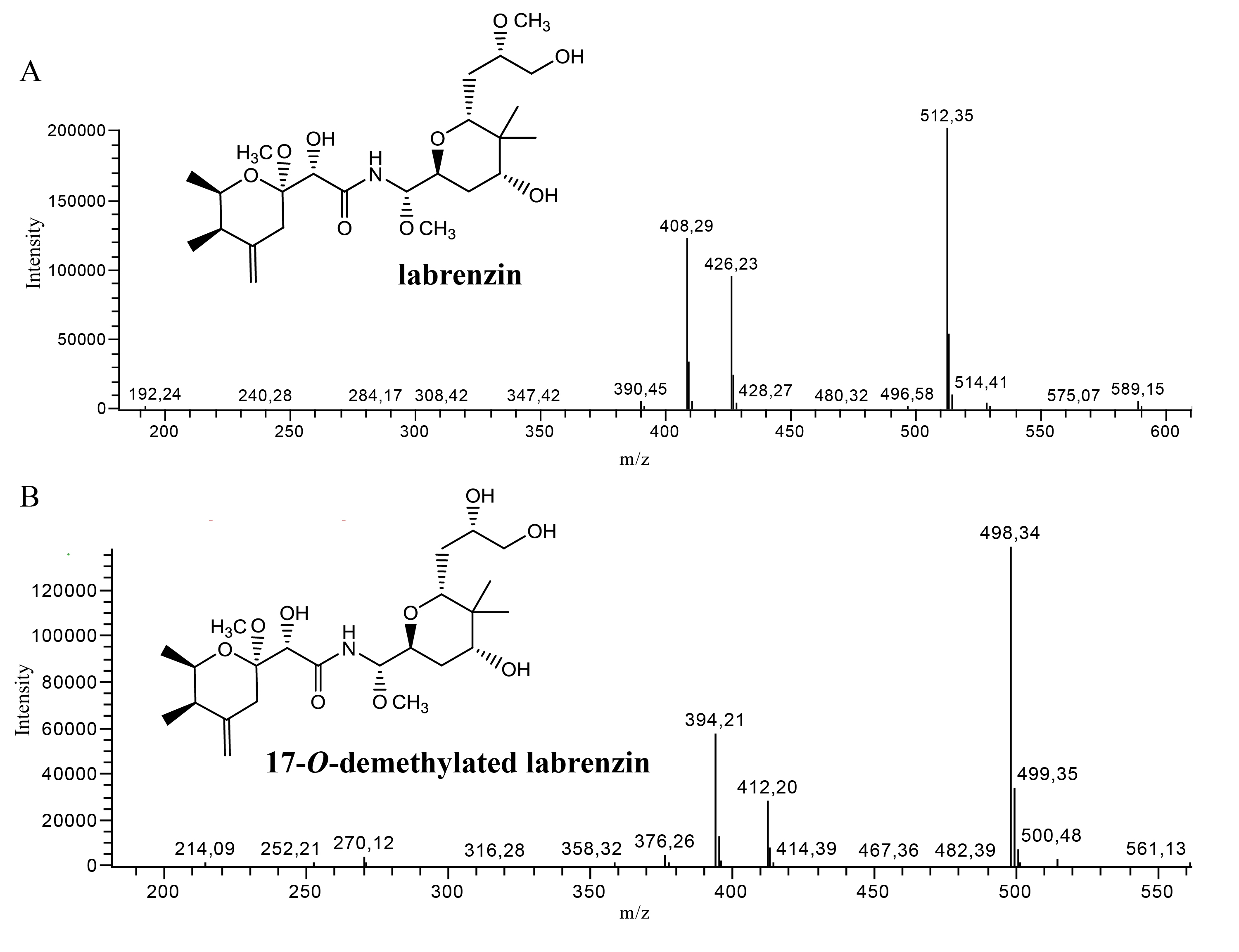
